# Early warning signals regarding environmental suitability in the *Drosophila* antenna

**DOI:** 10.1101/146522

**Authors:** Haoyang Rong, Prithwiraj Das, Adalee Lube, David Yang, Debajit Saha, Yehuda Ben-Shahar, Baranidharan Raman

## Abstract

**Highlights:**

- A novel geotaxis assay showed high intensity odorant exposures are harmful to flies
- Repulsion at high odor intensities can be a protective mechanism
- Olfactory receptor neuron (ORN) excitability abruptly changes with odor intensity
- A linear combination of ORN activities can robustly predict intensity-dependent behavioral repulsion

**Summary:** The olfactory system is uniquely positioned to warn an organism of environmental threats. Whether and how it encodes such information is not understood. Here, we examined this issue in the fruit fly *Drosophila melanogaster.* We found that intensity-dependent repulsion to chemicals safeguarded flies from harmful, high-intensity vapor exposures. To understand how sensory input changed as the odor valence switched from innocuous to threatening, we recorded from olfactory receptor neurons (ORNs) in the fly antenna. Primarily, we observed two response non-linearities: recruitment of non-active ORNs at higher intensities, and abrupt transitions in neural excitability from regular spiking to high-firing oscillatory regime. Although non-linearities observed in any single ORN was not a good indicator, a simple linear combination of firing events from multiple neurons provided robust recognition of threating/repulsive olfactory stimuli. In sum, our results reveal how information necessary to avoid environmental threats may also be encoded in the insect antenna.

## Introduction

Animals exhibit various degrees of behavioral preference to olfactory cues. They are attracted by food odors and pheromone, as a way to guarantee their survival and reproduction(Aron, 1979; Bronson, 1979; Reinhard et al., 2004; Wyatt, 2003). On the other hand, odorants produced by toxic substances that signal potential environmental danger lead to aversion(Stensmyr et al.; Zhang et al., 2005). While, odorants tend to maintain their overall odor valence over a wide range of concentrations, most drastically alter the polarity of odor valence as the concentration increases, i.e. the behavior preference switches from attraction to aversion(Stensmyr et al., 2003). What determines whether an odorant’s valence remains constant or changes with intensity?

The early olfactory circuits of *Drosophila* have been well studied both from an anatomical(Couto et al., 2005) and functional perspective(Hallem and Carlson, 2006). However, the rules that govern how sensory stimuli get translated to behavioral outcomes remains poorly understood. For example, changing the intensity of a stimulus is arguably the smallest manipulation to the sensory input possible as only the number of molecules is varied not its identity or other chemical features. Yet, existing behavioral data reveals that for many odorants the overall preference can switch as stimulus intensity is increased beyond a threshold value. This mismatch between the degree of variation in the sensory input and behavioral output raises the following important fundamental question: when and why do the same stimuli repel flies when delivered at higher intensities? And, how is this information encoded?

The repulsion to high intensity chemical vapors has been observed in many species(Poucher, 1974; Stensmyr et al., 2003; Yoshida et al., 2012), although, its significance is yet to be understood. Such a response is particularly confounding considering that many of these stimuli may otherwise not evoke any innate response, or even be attractive to them at lower intensities. How then are such stimuli represented in the olfactory system and what aspects of neural responses change abruptly with intensity? Electrophysiological and imaging studies have shown that increasing odor intensity activates additional olfactory receptor neurons that are not responding at a lower concentration(Duchamp-Viret et al., 2000; Knaden et al., 2012; Rubin and Katz, 1999; Semmelhack and Wang, 2009). Previous works have proposed that this recruitment of additional activity might be sufficient to explain both invariance in neural encoding(Asahina et al., 2009) and changes in behavioral response (i.e. ‘the recruitment hypothesis’)(Suh et al., 2004). If this recruitment hypothesis is indeed true, then behavioral variance with intensity may simply arise as a result of recruiting exclusive sensory channels or ‘labelled lines’ that mediate aversion(Knaden et al., 2012; Semmelhack and Wang, 2009). Whether such recruitment of additional activity at higher intensities happens for all or a subset of odorants is not clear. Furthermore, it is unclear whether the spiking activity in these additionally activated receptor neurons alone determines the odor intensity at which the behavioral preference to this stimulus switches from attraction to repulsion.

Here, we explored this issue in the *Drosophila* olfactory system. We found that flies were repelled by odorants at intensities beyond which the vapors were harmful to them. Exposure of flies to such high-intensity vapors anesthetized them. To understand how the information regarding odorants was encoded as their intensities were altered from innocuous to threatening, we recorded from olfactory receptor neurons on the antenna. Our results indicate that in addition to recruitment of receptor neurons at higher concentrations, abrupt transitions in neural excitability also occur as stimulus intensity is increased. Furthermore, our data reveals that while activity recruitment or excitability changes in receptor neurons may correlate with behavioral preference changes for some select odorants, they do not provide a general rule for translating sensory input to behavior. Notably, our results indicate that total spiking activity in a select few receptor neurons may serve as a robust indicator of changes in behavioral preference with intensity and thereby may act as a neural basis for an early warning signal.

## Results

### Behavioral switch to repulsion at high odor intensities

To identify general trends in dose-dependent behavioral preference changes, we used a stimulus set comprising of seven different odorants. Each stimulus was delivered over a wide concentration range (over seven log-units of magnitude) in order to include innocuous and threatening olfactory valences. It is worth noting that the stimulus set included many fruit-related odorants(Laissue and Vosshall, 2008) such as ethyl acetate, methyl acetate, ethyl butyrate, methyl hexanoate and ethyl-3-hydroxy butyrate. In addition, we included odorants such as 2,3-butanedione and 1-hexanol that are known to inhibit innate avoidance response due to CO_2_(Turner and Ray, 2009). We examined the behavioral preference of unstarved flies to each odorant on the panel using a standard T-Maze assay. We found that all seven odorants examined were non-repulsive at lower intensities, but the behavioral preference switched to strong aversion at higher intensities. The threshold concentration at which the preference switched varied a little between two subsets of odorants: 10^−2^ v/v or 10^−1^ v/v (n ≥ 8 for all concentrations but 10^−1^ concentrations for which n ≤ 5; two-tailed one-sample *t*-test, p values<0.05; Figure 1A). Nevertheless, our behavioral data suggests that the switch in overall behavioral preference with intensity may be a common feature in this sensory modality.

**Figure 1:**
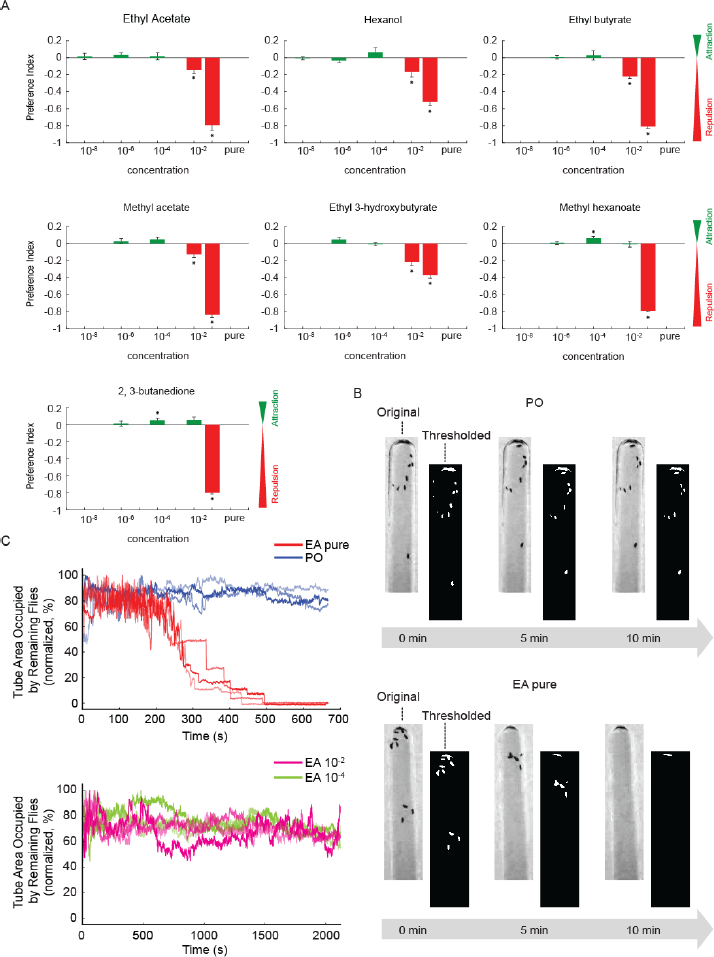
Dose-dependent behavioral responses to odorants. (A) Behavior preferences of fruit flies to different odor-intensity combinations were assayed using a T-maze assay and shown. Positive and negative preference index values represents attraction and repulsion respectively. Mean ± s.e.m is shown for all concentrations (n ≥ 8 for all concentrations but 10^−1^ concentrations for which n ≤ 5). Asterisks indicate significant increase or decrease in behavioral preference values at p<0.05. (B) Representative results from a geotaxis assay are shown. Note that while the flies clung onto the test tube walls they were also exposed to either paraffin oil vapors (control; top panel) or ethyl acetate vapors (bottom panel). Both original and the thresholded image highlighting the position of flies (in white) on the test tube walls are shown for three different time points. Note that the number of flies stuck to the walls reduced over time when they were exposed to high-intensity ethyl acetate vapors. (C) The area of test tube wall occupied by flies (y-axis) was tracked as a function of time and plotted for four different conditions: paraffin oil (PO; blue traces in the top panel; n=3), undiluted ethyl acetate vapors (EA; red traces in the top panel; n=3), ethyl acetate at 10^−2^ (magenta traces in the bottom panel; n=3), and 10^−4^ ethyl acetate vapors (green traces in the bottom panel; n=3). Each curve was normalized by its maximum to facilitate comparison across experiments. Note that the area occupied by flies on the tube walls dropped to zero only for all pure ethyl acetate cases.

Next, we sought to understand the need for repulsion at higher odor intensities. We found that most flies exposed to odorants beyond the repulsion intensities were anesthetized before they could make a decision to enter a T-maze arm. To quantitatively illustrate this, we used another behavioral assay where flies performing geotaxis were exposed to high-intensity vapors of ethyl acetate. We found such high-intensity vapor exposures were unsuitable to flies, and those performing geotaxis were anesthetized and fell from the walls of the climbing tubes. Whereas, control exposures to paraffin oil had no such effect on flies and they managed to hang onto the walls for the entire duration of the experiment.

We tracked the area of the climbing tube that was occupied by the flies as a function of time. As can be expected, this metric remained stable for flies exposed to the paraffin oil, but reduced to zero for high-intensity ethyl acetate exposures (n=3 assays for each condition, Figure 1B **and** C)). This effect of high-intensity ethyl acetate vapors on flies was not observed during low intensity exposures (10^−4^ v/v and 10^−2^ v/v) of the same odorant. It might be worth to note that 10^−2^ v/v was the threshold intensity when the overall behavioral preference switched to repulsion for this odorant. Taken together, these results indicate that repulsive response of flies to high-intensity chemical vapors is a protective mechanism that allows them to avoid exposures to harmful chemicals.

### Olfactory receptor neurons’ response non-linearities

Given the drastic change in the behavioral response for all odorants tested, we examined how the sensory input from ORNs change with odor intensity. First, we performed extracellular recordings from fruit fly ORNs in the ab3 sensillum when the antenna was puffed with ethyl acetate vapors at different concentrations (schematically shown in Figure 2A). We found that both neurons in the ab3 sensillum were not activated at low intensities of ethyl acetate exposure but became activated at a threshold concentration of 10^−2^ v/v (Figure 2B). Since the ab3 neurons were recruited at a certain threshold intensity of the odorant, we examined if this recruitment correlated with the behavioral preference switch. As can be noted, the increase in neural activity in this sensory channel (n=10 sensilla, paired *t*-test, Bonferroni corrected p< 0.0125) reflects when flies were repelled by ethyl acetate in the T-Maze assay (Figure 2C). Therefore, these neural and behavioral data taken together suggest that recruitment of spiking activities in additional receptor neurons may correlate with intensity-dependent behavioral response switch for ethyl acetate.

**Figure 2.**
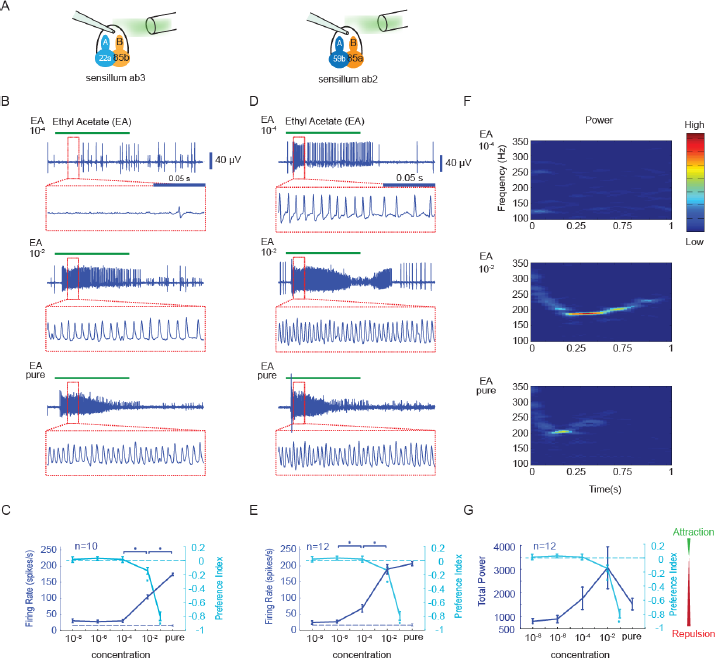
Recruitment vs. abrupt transitions in receptor neuron spiking. (A) A schematic of ab2 and ab3 sensillum recordings is shown. (B) Representative extracellular recording traces acquired from a ab3 sensillum are shown. Raw traces were high-pass filtered at 55 Hz to remove the DC-component. Responses elicited by ethyl acetate vapors delivered at 10^−4^, 10^−2^ and undiluted intensities are shown. The green bar above the voltage traces indicate the 1 s time window when the stimulus was presented. A small 150 ms segment (red boxes) of the recording was magnified and shown underneath each raw trace for clarity. (C) The total spiking activity of neurons housed in the ab3 sensillum is plotted as a function of stimulus concentrations (blue, mean±s.e.m.). The dotted line (at the bottom of plot) indicates the spiking activities elicited by paraffin oil exposures. Asterisks indicate significant increase of firing rate compared with the neighboring lower concentration (p<0.05, paired *t*-tests, n=10 trials). For comparison, the behavioral preference index (cyan, mean±s.e.m.) observed in the T-maze assay for various intensities of ethyl acetate exposures is also shown. Note that the behavioral preference switched to repulsion at 10^−2^ ethyl acetate exposures. (D) Same as 2b, but showing responses of ab2 neurons to ethyl acetate at various intensities. Note that ethyl acetate exposures clearly elicit a detectable response at 10^−4^ dilution. However, note that at higher ethyl acetate intensities only oscillatory field potentials of varying amplitudes are observed. (E) Similar comparison as in panel c, but comparision between firing rates of neurons in ab2 sensilla (blue, mean±s.e.m.) with the behavioral preference (cyan, mean±s.e.m.) is shown for ethyl acetate presented at various concentrations. (F) A moving window power spectra of a representative extracellular trace recorded from ab2 sensillum is shown. Power in the high gamma band frequencies (>150Hz) can be observed during 10^−2^ and undiluted ethyl acetate exposures. (G) Similar comparison as in panel c, but comparison between the total power of signals from ab2 sensilla (blue, mean±s.e.m.) and the behavioral preference (cyan, mean±s.e.m.) is shown for ethyl acetate presented at various concentrations.

Next, we examined how neural activities in other receptor neurons that were strongly activated by ethyl acetate (Or59b expressing ORN housed in the ab2 sensillum) were altered as a function of stimulus intensity (Figure 2A). Consistent with existing data(Hallem and Carlson, 2006), we found that at lower intensities the spiking activity increased beyond the baseline levels particularly for the ab2A neuron expressing Or59b receptor (Figure 2D). However, as the odor intensity was increased beyond a threshold concentration (10^−2^ for ethyl acetate) the spiking activity transitioned from clearly distinguishable spikes to a response regime where individual action potentials were no longer resolvable (Figure 2D). Rather, it appeared that spikes collided with each other and generated oscillatory field potential activity with increased power in the high-gamma band (~ 200 Hz; Figure 2F). This oscillatory extracellular activity was detected in all our ab2 sensilla extracellular recordings following exposures to high concentrations of this odorant (n=12 sensilla). Notably, both the frequency content of the field potential activity and its amplitude varied as a function of ethyl acetate intensity (Figure 2D).

Since we were unable to resolve individual spikes at high intensities, to characterize the dose-response curve, we counted the total number of firing events during any single ethyl acetate exposure and plotted it as a function of odor intensity (n=12 sensilla, paired *t*-test, Bonferroni corrected p< 0.0125; Figure 2E; see Methods). The mean dose-response curve was sigmoidal with the number of firing events making an abrupt increase right when the extracellular activity transitioned from spiking to oscillatory field potentials. Interestingly, a qualitatively similar dose-response curve could also be generated by examining the total change in oscillator power in the high-gamma range (Figure 2F **and** G). More importantly, the switch in behavioral preference for ethyl acetate occurred right at the threshold intensity when the neural activity in the ab2 neurons switched.

These results, taken together suggest that both recruitment of additional receptor neurons’ activities and an abrupt switch in receptor neuron firing pattern (from low to high) may both correspond to the switch in the overall behavioral preference for ethyl acetate at higher intensities.

### Field potential oscillations in olfactory sensillum

How are the receptor neuron oscillations generated? To understand this issue, we added Na+ channel antagonist tetrodotoxin (TTX) to recording glass pipettes(Nagel and Wilson, 2011). This pharmacological manipulation resulted in elimination of all ORN spiking activity and also abolished field potential oscillations observed at high intensities (Figure 3A). However, note that the DC-component of the sensilla local field potential caused by transduction currents remained unaffected. These results confirm that the field potential oscillations are not an artifact of our extracellular recording approach as they can be abolished using Na+ channel blocker. Furthermore, note that the DC component of the signal is monotonic with odor intensity (Figure 3A). In sum these results suggest that the oscillatory potentials must originate downstream of the transduction machinery possibly due to collision of spikes.

**Figure 3:**
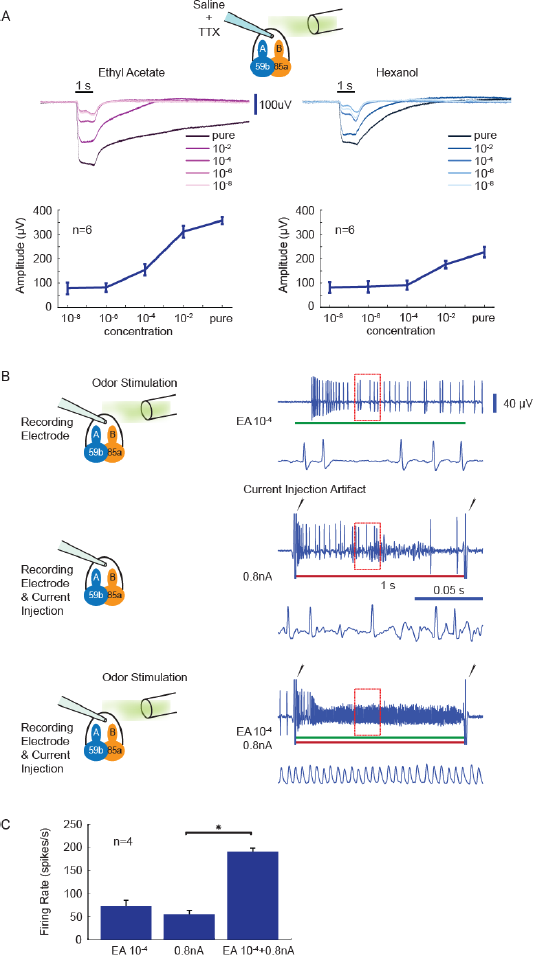
Mechanism underlying field potential oscillations in olfactory sensillum. (A) Extracellular recordings obtained from an ab2 sensillum with a glass pipette filled with saline and tetrodotoxin (TTX) is shown (see methods). TTX blocked all sodium spikes but the transduction potential due to activation of olfactory receptors by ethyl acetate (left) of hexanol (right) was still observed. The amplitudes of the transduction potential (i.e. magnitude of the DC component) increased monotonically with the concentration for both odorants. Note that neither spikes nor oscillatory field potentials can be observed. Bottom panel: average amplitude of the DC component from 6 trials plotted as a function of stimulus intensities is shown for both ethyl acetate and hexanol. (B) Extracellular recordings obtained from an ab2 sensillum are shown for three different cases: (top row) exposure to 10^−4^ ethyl acetate (second row), direct current injection (0.8 nA) into the sensillum, and (third row) a simultaneous presentation of both ethyl acetate at 10^−4^and current injection (0.8 nA). The color bars at the bottom of the trace indicate when the odor puff and/or current injections were delivered. Black arrows indicate stimulation artifacts at the onset and offset of current injection. Note that neither odor stimulation (EA at 10^−4^), nor current injection alone could generate oscillatory extracellular field potentials. However, when they were presented together, oscillations could be observed. (C) Bar plot quantifying the ab2 firing rates observed during the three conditions presented in **panel b.**

To test the spike collision hypothesis, we examined whether this transition from low firing spiking regime to a high-firing one could be controlled by pairing odor stimulation with electrical stimulation. As noted previously, ethyl acetate at 10^−4^ dilution elicited clearly resolvable spikes. Similarly, a weak current injection (0.8 nA) alone generated modest increase in spiking activity in the receptor neurons housed in ab2 sensillum. However, when the odor stimulation was combined with the current injection, we found that the extracellular activity transitioned to the oscillatory field potential very similar to those observed at high odor intensities (Figure 3B). These results taken together with the pharmacological manipulation findings confirm that the non-linear switch to a high firing oscillatory field potential regime in ab2 receptor neurons is due to modulation of excitability in these neurons.

Could the recently identified non-synaptic interactions between receptor neurons(Su et al., 2012) influence encoding of stimulus intensity? To examine this issue, we generated transgenic flies with only one functional receptor neuron in the ab2 sensilla. We examined the responses of these transgenic flies to ethyl acetate and compared the same with those obtained from wild-type flies (Figure 4A). Note that transgenic flies with genetically ablated Or85a (n=6 sensilla) or Or59b (n= 7 sensilla) expressing receptor neuron still reveal similar transitions in spiking activity with increase in stimulus intensity (paired t-test, Bonferroni corrected p< 0.0125; Figure 4A **and** B). Hence, we conclude that interactions between these receptor neurons are not necessary to mediate the observed modulation in their excitability.

**Figure 4:**
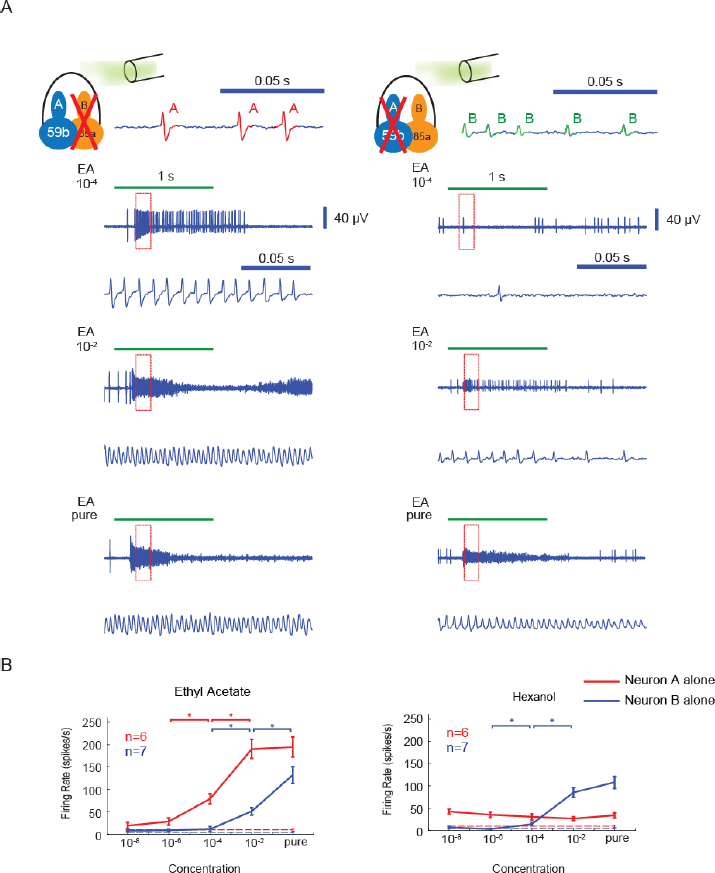
Individual receptor neurons can generate oscillatory field potentials. (A) Sensillum recording obtained from transgenic flies with either the B neuron (Or85a) or A neuron (Or59b) in the ab2 sensillum ablated are shown. Top: Schematics showing ORN ablation and actual extracellular trace obtained from such genetically modified sensillum are shown. Note only spikes of single amplitude are observed after ablation of one receptor neuron. Bottom panels: Representative extracellular recording traces showing responses elicited by ethyl acetate at different intensities are shown. Note that oscillatory field potentials could still be observed at high ethyl acetate intensities when only A neuron or B neuron remained. (B) EA and Hex Dose-response curves for ab2A and ab2B neurons are shown. Asterisks indicate significant increase of firing rate compared with the neighboring lower concentration (p<0.05, paired t-tests).

### Oscillatory dynamics in a Hodgkin-Huxley type neuron

To understand how the same neuron can create firing events of varying shapes, we performed a phase plane analysis of neural excitability for a Hodgkin-Huxley type neuron model (HH model). To perform this 2-d analysis, we reduced the HH model from a system of four ordinary differential equations to two by making two assumptions (see Methods). First, we assumed that the sodium channel activation gating variable *‘m’* reaches its asymptotic value instantaneously to eliminate one variable. Second, by expressing the sodium channel inactivation gating variable *‘h’* as a linear function of the potassium channel activation gating variable *‘n’* we eliminated the second variable.

We found that this reduced HH model could generate spiking activities of different shapes depending on the amplitude of the input current. For depolarizing input up to a certain threshold value, we were able to observe clearly resolvable tri-phasic individual action potential waveforms (Figure 5). Beyond the threshold value, we found the action potential waveform shapes became considerably narrower with smaller peak to trough amplitudes. The spikes produced appeared qualitatively similar to the oscillatory extracellular potentials observed in ORNs during high-intensity odor exposures.

**Figure 5:**
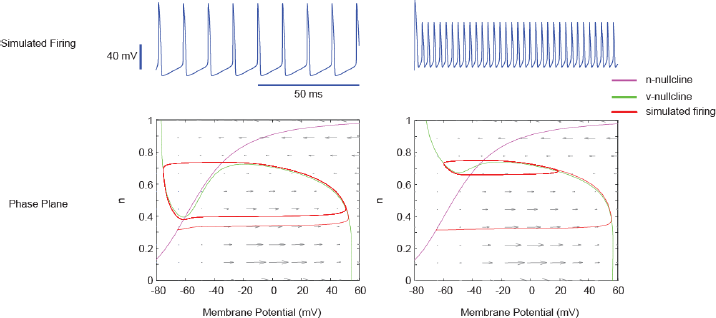
Modulation of limit cycle size and shapes in a Hodgkin-Huxley model. Simulated spikes of different amplitudes are shown from a 2-dimensional reduction of Hodgkin-Huxley model are shown (see Methods). All parameters in the model were kept constant for both cases but the amplitude of injected current was substantially increased from 10 μA (left panel) to 175 μA (right panel). **Phase-plane analysis:** The fast variable (membrane potential; x-axis) is plotted against slow variable (potassium gating variable) for the two different spiking conditions shown above. The fast and slow variable null clines (curves along which the derivatives are zeros) are shown in green and purple, respectively. Gray arrows indicate the direction the system would evolve in the locality of a specific region. The red trace illustrates the simulated firing evolved in the phase plane. Notice that the shape of cubic v-nullcline changes substantially for high current injections thereby making the limit cycle almost bi-phasic.

We found that this change in action potential shape was mainly due to the alterations in the dynamics of the fast variable (corresponds to the membrane potential of the neurons, ‘V’ in the HH model). Note that the shape of the fast variable *null cline* (the curve along which the membrane potential is held constant) changed depending on the magnitude of the depolarizing current input. This resulted in the shape of the period events changed drastically (i.e. limit cycle in dynamical systems jargon, or, action potentials fired by the neuron model, its biological interpretation). Therefore, these results further support our interpretation that the changes in the spiking activity observed in our receptor neuron recordings could arise due to the changes in ORN excitability.

### Rules for predicting behavioral preference changes

Finally, we examined how well spiking activities in receptor neurons (paired *t*-test, Bonferroni corrected p< 0.0125; Figure S1) correlated with the behavioral preference switch for all odorants in the panel. Our results indicate that spiking activities in ab2 or ab3 receptor neurons when considered individually correlated with the behavioral preference switch in only a select few odorants (Figure 6A). This result was confirmed by plotting the ab2 and ab3 neural spiking response versus behavioral preference for a given odorant at a particular intensity (Figure 6A). A single threshold that separates non-repulsive stimuli from repellent odor-intensity combinations could not be found. However, when we linearly combined the contribution of both these sensory channels, we found that the total activity in these two sensory channels could robustly identify which odorants at what concentrations evoked an attractive or a repulsive response (Figure 6B).

**Figure 6:**
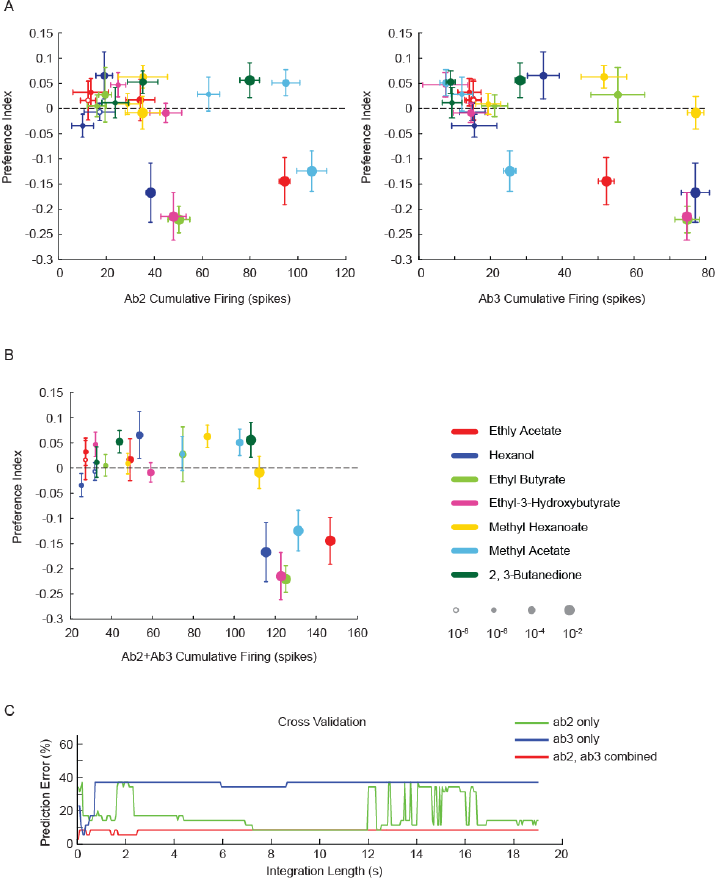
Predicting behavioral preference from receptor neuron responses. (A) Behavioral preference for each odorant at each intensity (y-axis) is plotted against cumulative spike counts (500 ms integration window since response onset) is shown for both ab2 (left panel) and ab3 (right panel) sensillum. The mean±s.e.m for spiking activity and the behavioral preference values for each stimulus used in the study are shown. The size, fill and color of the marker uniquely identify odor identity – intensity combination. In both panels, note that a single threshold firing rate that reliably separated repulsive odors from non-repulsive ones did not exist. (B) Behavioral preference plotted against the sum of cumulative firing (500ms) from both ab2 and ab3 sensilla are shown. Note that stimuli that evoke less than 110 cumulative spikes/s in these two channels were non-repulsive, whereas those odor-intensity combination that evoked more than this threshold repelled flies strongly. (C) Performance characterization of an optimal linear classifier is shown for three different cases: (i) using spiking information from ab2 sensilum alone (green) (ii) using spiking information from ab3 sensillim alone (blue), and (iii) using information from both these channels (red). The prediction error is shown for different values of the integration window used to summate the spikes. A leave-one-odor-out cross-validation scheme was used to quantify performance in this plot.

The analyses presented so far examined the segregation of behavioral preferences based on spike counts from two sensory channels but within a specific time window (500 ms from odor onset). How robust are these results when this assumption regarding the spike integration window is removed? To examine this, we compared the prediction performance of an optimal classifier (linear SVM) when the classifier was trained using information obtained from either a single channel (ab2 or ab3; 1-d problem), or from both channels (ab2 and ab3; 2-D problem). We characterized the prediction error for these three cases as a function of spike integration window length (Figure 6C).

We found that the total spiking activities in the *ab3* sensory hair alone could provide low prediction error when the integration window was set to a specific value (250 ms). However, beyond this value, the prediction error increased significantly. On the other hand, the total spikes from *ab2* sensory hair supported predictions with higher error rates for a wide range of integration window durations. Neither of these two sensory channels, when considered in isolation could support rapid decision making (< 100 ms) with low prediction error. However, the prediction error when spiking responses from both *ab2* and *ab3* channels were simultaneously considered, the prediction error became less sensitive to the integration window length. Furthermore, as can be expected, the combinatorial approach could achieve the lowest prediction error among all three cases within the first 50 ms after response onset. Since flies are capable of making decisions rapidly (within 100 ms)(Bhandawat et al., 2010; Steck et al., 2012), as might be needed for an escape response, these results further support the need for a readout scheme based on spiking information from multiple sensory channels.

Taken together, our result suggests that a perceptron-like “summation and thresholding” model, in which a linear combination of information from multiple ORN types can robustly explain the behavioral response.

### Discussion

The volatile nature of chemosensory cues transduced by the olfactory system indicates that it is well suited to serve as a first responder capable of informing an organism about potential environmental hazards. However, to generate an early warning signal two additional requirement need to be considered. First, given that the set of chemical stimuli that are harmful to an organism may be broad, a more general encoding strategy (many inputs–to–one behavioral outcome) may be required. Second, the sensory cues must be mapped onto a behavioral response that will help avoid such threats (i.e. repulsion). Our results indicate that the *Drosophila* olfactory system does indeed use a general strategy based on total spikes from multiple sensory channels to encode such information. Furthermore, dose-dependent odor-evoked repulsion observed in many organisms including fruit flies may help avoid such environmental threats.

We found that exposures to most volatile organic chemicals beyond a certain threshold intensity repelled fruit flies. This was true even for those considered to be food odorants(Laissue and Vosshall, 2008; Stensmyr et al., 2003). Exposures beyond this threshold were unviable to flies, and those performing geotaxis during such exposures were anesthetized and fell from the walls of the climbing tubes. To understand how the information regarding odorants were encoded as their intensities was altered from innocuous to threatening for flies, we recorded from olfactory receptor neurons housed in large basiconic sensory hairs on the antenna. Our results indicate that when information from select few receptor neurons in ab2 and ab3 sensillum were combined, we could robustly predict which odorant at what intensity became repellant and therefore not suitable for flies.

To test the robustness of our results and conclusions, we performed a similar classification analyses but using another published dataset(Hallem and Carlson, 2006). We compared the performances of three different integration strategies to predict the behavioral outcome: (i) combine inputs from all receptor neurons irrespective of which type of sensory hair they are housed in (Figure S2A), (ii) integrating spiking responses of all antennal basiconic type ORNs (Figure S2B), and (iii) combining signals from select neurons in ab2 and ab3 sensillum (Figure S2C; similar to the analysis presented in Figure 6). As a general rule, we found that the classification performance increased monotonically with the number of pooled ORN types (Figure S2). However, when the antennal basiconic neurons were exclusively combined classification performance increased much faster and reached higher asymptotic success rates than the ‘all ORN strategy’ (Figure S2B vs. Figure S2A). Alternately, when select neurons in ab2 and ab3 sensilla were integrated, again good discrimination between innocuous and repulsive cues were observed. These results further provide an independent corroboration of our findings and suggest that either integration from a sub-type of receptor neurons (i.e. all housed in basiconic type sensory hairs), or from a select few sensory channels provide effective approaches to translate sensory inputs into behavioral outputs.

Previous work on odor-evoked repulsion in flies either using stress-related odorants(Suh et al., 2004) or unsuitable food sources(Stensmyr et al.) suggested a labelled line approach for the transformation of sensory input onto an avoidance response. In this work, our results suggest a combinatorial approach for generating the same motor response. Although these results may potentially be seen at odds with each other, it is quite possible that multiple mapping schemes from stimulus space to behavior could co-exist. Alternately, the combinatorial input from receptor neurons may be transformed to activate labelled lines in the downstream neural circuits that could then evoke repulsion.

These sensory-motor transformations could alternately be viewed from the perspective of metabolic costs. Since spiking is metabolically expensive(Laughlin et al., 1998), the increase in total spiking activities indicate an expensive operation. A previous study using Drosophila larvae found that odorants that evoked more inhibition were also more likely to be repulsive(Kreher et al., 2008). Therefore, a sigmoidal sensory-motor transformation that maps too much or too few spiking (extremes of metabolic costs) onto repulsion seems to account for results reported here by us and elsewhere by others(Kreher et al., 2008). Whether this result is merely correlational or is metabolic costs an important variable that can shape behavioral outcomes needs to be systematically determined.

Finally, we found that at extremely high stimulus intensities, the clearly resolvable spiking activity in individual neurons transformed into oscillatory field potential activity with power in the high-gamma frequencies. We found that this abrupt transition in spiking behavior is largely due to changes in neural excitability and can be abolished with TTX or induced with current injections. Further, such oscillatory activities can be observed when multiple cues, which by themselves do not generate such a response, are combined. Indeed, we found that olfactory mixtures reported in another pioneering study on non-synaptic inhibition between co-housed receptor neurons did indeed evoke oscillatory field potentials of varying amplitudes similar to those reported here. Therefore, we conclude that complex changes in spiking behavior of receptor neurons can simply be induced due neural excitability modulations and without any coupling between them.

What then might be the need for such high activity regimes, given the strong synapses between the receptor neurons and their downstream targets in the antennal lobe(Wilson, 2013), and the recent report that behavioral response can be generated with modest number of spikes(Bell and Wilson, 2016)? It is possible that such responses may be an unavoidable consequence of having high sensitivity to food-related odorants. While responses to extremely low concentrations may guarantee sustenance, a compensatory mechanism might be needed to avoid the same odorants when they become unsuitable.

## Methods

### Fly Stocks

Flies (Canton-S) were raised on cornmeal medium at 25 ± 1°C under 12:12 light-dark cycle. For experiments with transgenic flies (Figure 4), ORNs were selectively ablated by crossing UAS-DTI flies with Or59b-Gal4 or Or85a-Gal4 lines.

### Odor Stimuli

7 odorants were used in both electrophysiology and behavior experiments: 2,3-butanedione (97%, Sigma-Aldrich Co. LLC.), ethyl acetate (99.8%, Sigma-Aldrich Co. LLC.), ethyl butyrate (99%, Sigma-Aldrich Co. LLC.), ethyl-3-hydroxybutyrate (≥97%, SAFC, Sigma-Aldrich Co. LLC.), hexanol (≥98%, SAFC, Sigma-Aldrich Co. LLC.), methyl acetate (≥99%, SAFC, Sigma-Aldrich Co. LLC.), and methyl hexanoate (≥99%, SAFC, Sigma-Aldrich Co. LLC.). Except pure odors, all dilutions were made by dissolving pure odor solutions in paraffin oil (J.T. Baker).

### Single-Sensillum Recordings

Female flies aged from 5-8 days after eclosion were used. To perform extracellular recordings from receptor neurons we followed a previously published procedure(Dobritsa et al., 2003). The fly antenna was extended and fixed using a glass capillary on a coverslip. To acquire action potentials, a glass electrode filled with saline (impedance ~40MΩ) was inserted into the middle portion of a sensillum. Another reference glass electrode was inserted into the contralateral eye. The signals were amplified (gain =10; Axon 900A, Molecular Devices) and filtered with a high-pass filter set to DC and a low-pass filter set at 10kHz. A custom Labview software was used to acquire samples at 15 kHz. No more than two sensilla from the same fly were recorded.

For each of the 7 odors described above, we tested 5 concentrations: undiluted, 10^−2^, 10^-4^, 10^−6^, and 10^−8^dilutions.

For each trial, 50μL odor dilution was added to a filter paper strip and placed in a Pasteur pipette(Dobritsa et al., 2003). A humidified, carrier air stream at a flowrate of 2000 sccm was directed at the fly antenna throughout the experiment. To present an odor stimulus, a 200 sccm air puff was passed through the filter paper strip containing the odor solution and into the carrier airstream.

Each trial lasted 60s with an intertrial interval > 30s, and the stimulus was delivered from the 10^th^ second to the 11^th^ second of the trial. Odors were presented in pseudo-random blocks based on odor identity. Different concentrations of a single odorant were also presented in a random order, except for the undiluted stimuli which was always presented as the last stimulus in each block.

### T-maze Behavior Assay

We used 5 – 8 day old male and female flies. To be consistent with electrophysiology experiments, flies used in our behavioral experiments were also unstarved.

We tested the same 7 odors used in our electrophysiology experiments. Each odorant was presented at the following concentrations: 10^−1^, 10^−2^, 10^−4^, 10^−6^, and 10^−8^ v/v.

One piece of folded filter paper was placed at the end of each of the two plastic test tubes (17mm×100mm, 14mL Round-Bottom Polypropylene Tubes, Falcon). After adding 50μL odor dilution and paraffin oil to the filter papers in each of the two test tubes, they were sealed with Paraflim (Bemis Company, Inc.). To allow sufficient evaporation of odorants, test tubes were left undisturbed for ~10mins before further use. For each trial, 150~200 flies were placed into the T-maze fly chamber. Assays were conducted in a dark room to prevent interference from any visual cues. Before testing, flies were given 1 min acclimatization time. Then, the fly chamber was lowered to allow the flies to access the two test tube arms. The flies were given 1 min to make their decision. The preference index was calculated using the following formula:

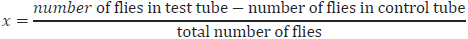

### Geotaxis Behavior Assay

Unstarved flies aged between 5-9 days were used. 8-12 flies were placed in test tubes (17mm×100mm, 14mL Round-Bottom Polypropylene Tubes, Falcon). To prevent flies from escaping the test tube, a piece of metal mesh was attached to cover the open end of the test tube. The tube was inverted and kept perpendicular to the table, so the flies could climb toward the top of the tube. Another test tube was cut off about 1.7 cm from the opening, and one end sealed using a round glass cover-slip (12-546-2, Fisherbrand) to form a manifold for a filter paper containing the odor solution. Right before the experiment, 50μL of odor dilution or paraffin oil was added to the filter paper and placed in this manifold. The test tube with flies was inverted on top of the odor manifold. The connection between the tubes was sealed with dental wax (Surgident, Heraeus Kulzer Inc.). The assembly was then placed in front of a red LED panel. Fly movements on the test tube walls were recorded using a camera (C920, Logitech) at 30 frames/s.

Movies were analyzed using OpenCV 3.0.0 in Python 2.7.10. Region of interest (ROI) was manually picked to track the fly movement. The starting time for each trial was manually set to be the frame at which the assembly was stably placed in front of the background light panel and the camera self-adjusted to a stable setting. Only signals from the blue channel were used, so the frames became gray-scale with each pixel value ranging from 0 to 255. Every frame was thresholded to separate shadows created by flies from those of the test tube itself. Number of pixels with a value below a threshold value (set to 100) was regarded as the total shaded area. To obtain the fly occupancy area (FOA) in each frame, the tube shaded area was subtracted from the total shade area. To compute the tube shaded area, we averaged across all 20000 frames from each movie to obtain the average frame. In the average frame, the shade created by the tube itself was much easier to be differentiated from the ones created by flies. Plot shown in Figure1c was generated by passing area occupied by flies across frames through a 30-point moving average filter and normalized to the maximum of that curve.

### Pharmacology

Tetrodotoxin (Sigma-Aldrich Co. LLC.) was dissolved in Ringer solution to a concentration of 50μM/L. The extracellular recording electrode was filled with the diluted TTX solution and inserted into the sensillum. About 10~15 min was given to allow the perfusion of TTX and abolish sodium spikes. The recording was not performed until spontaneous spikes were no longer observable, which was visually checked on an oscilloscope.

### Current injection

Positive current 0.8nA was injected into the sensillum through saline filled glass electrode. ORN activities were recorded for current injection, a low concentration of ethyl acetate, and a combined stimulus of current (0.8nA) and odorant (ethyl acetate) presented simultaneously to the antenna. The current injection and odor delivery were 1s in duration.

### Determination of response onset

We observed that since the odor puff had to travel a distance before reaching the antenna, ORN response onsets occurred after varying delays following stimulus onset across experiments. Therefore, to precisely determine the ORN response onset, we used a metric based on changes in field potential recorded from the sensillum. More precisely, we computed the first derivative of the band-pass filtered baseline (2nd order Butterworth band-pass filter, 0.1~5 Hz). The first time point after stimulus onset when the field potential’s derivative exceeds a chosen threshold was treated as the time of ORN response onset.

For most traces, their response onset was directly decided by its baseline drop. For traces without a detectable baseline drop at low odor intensities, their response onset was determined by the average response onsets of the same odor at higher concentrations (pure, 10^−2^).

### Firing event detection

Firing events were detected by a custom routine which in principle detected voltage peaks above a preset threshold (usually 4.5 times the pre-stimulus baseline s.d.).

When ORN responses enter high-activity levels (typically > 200 Hz), low-amplitude oscillatiory waveforms (LAOs) were observed. Therefore LAOs were extremely difficult to detect using the thresholding method. To address this issue, we developed a template-matching algorithm. In this algorithm, signal segments were binned in a short moving window that was compared with an oscillation waveform template. If the signal segment in a particular moving window was similar enough to the template, then the signal segment was counted as an LAO.

To create a template for oscillation waveform, a trace segment with typical, consistent oscillations was manually selected. In this segment, each oscillatory event could be robustly detected due to their large amplitudes. These oscillatory events were peak aligned and binned so that each bin solely contained the complete waveform of only one oscillatory event. Binned waveforms were each normalized by subtracting the mean and dividing by the standard deviation of signals of each time bin. The normalized waveforms were then averaged over 1875 such normalized oscillatory events to generate a oscillation waveform template.

To clean up the original trace for template matching, detectable supra-threshold firing events, including spikes and oscillations, were first removed. The remaining trace was concatenated and binned into 50 ms non-overlapping time segments. Power was computed for each 50 ms time segment. Consecutive segments with power larger than a preselected threshold were considered to contain LAOs. These LAOs-containing bins were again concatenated and pattern matched with the oscillation waveform template. Signals in the moving window were normalized as described above. The angular distance between the windowed signal (V_s_) and template (V_t_) was calculated to quantify their similarity. Because the window moved by one data point every step, we could obtain a trace of angular distance with high temporal resolution. The local peaks in the angular distance trace with a value > 0.7 were considered to indicate LAOs.

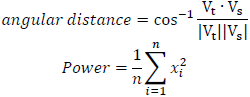

where *χ_i_* is the extracellular voltage recorded at time point *i*, and n is the total number of time points within a time bin.

### Classification analysis

We used a linear, optimal margin classifier- support vector machine (SVM) to predict the behavioral outcome (repulsion or non-repulsion based on T-maze results) given the spike counts from a combination of receptor neurons (present in ab2 or ab3 or both). The length of the window used to compute odor-evoked spike counts was systematically varied to quantify performance for different integration length (50 ms to 20 s). A soft margin version of SVM was used to make it more resistant to outliers. A leave-one-out cross-validation scheme (neural and behavioral data for one odorant at one intensity was left out; 34 odor-intensity combinations for training and 1 odorant-intensity for testing) to quantify our results.

Note since flies passed out before they could make a decision when exposed to pure odorants, they were regarded repulsive for the purposes of this analysis.

When only considering spike counts from the two neurons housed in a single sensillum type (i.e. ab2 alone or ab3 alone), we made predictions based on thresholding the input (i.e. if the input is above the threshold, the odor to be repulsive). The threshold value that resulted in the lowest training error was used. If multiple thresholds generated similar training errors, then the threshold that divides the data more evenly was picked.

We further tested our hypothesis using a published dataset(Hallem and Carlson, 2006), which contained mean firing rate of 24 types of ORNs to ten odors at four concentrations (10^−2^, 10^−4^, 10^−6^, and 10^−8^). We found four odors which are also in our research: ethyl acetate, hexanol, ethyl butyrate and 2,3-butanedione. The calculation of success rate was formulated as an “n choose k” problem, where n denotes the total number of ORN types available, and k denotes the number of pooled ORNs. Within each combination, we summed the firing rates of all k pooled ORNs and threshold the sums. If there existed a threshold that could correctly predict the odor valence, we counted it as a “success". If the total number of combinations and “successes” in a given “n choose k” problem was A and S, respectively. we calculated success rate as S/A.

### Modeling of Spikes and Oscillation

We simulated regular spikes and oscillation using a reduced two dimensional Hodgkin-Huxley model (HH model) derived from the standard HH model.

The standard HH model describes the membrane potential of a neuron with a set of four ordinary differential equations (ODEs):

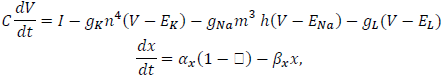
where *χ* can be replaced by n, m, and h. V stands for membrane potential. m, n, and h are gating variables with values in the [0,1] range. C is the membrane capacitance. *g_K_*, *g_Na_*, and *g_L_* are the maximum conductance of potassium, sodium and leak channels, respectively. represents reversal potentials of corresponding channels (E_Na_ = 55 mV, E_k_ = −77 mV and E_L_ = 61 mV).

Dimensionality reduced HH model: h can be replaced by a linear function of n, since n+h is almost a constant. m can be approximated by a simple polynomial equation. Thus, the standard HH model can be effectively approximated by a two-dimensional version of the model:

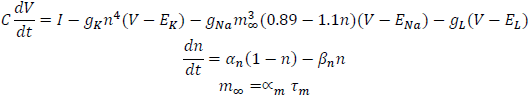

We set constant value for *I* to make it a constant stimulus. The membrane potential changes caused by a certain amplitude of input can be obtained by solving the ODEs. To simulate regular spiking activities and oscillations, we set *I* to equal 10 μA and 175 μA, and obtained firing rates of ~90 Hz and ~340 Hz respectively.

### Statistical Tests

No statistical method was used to predetermine the sample size.

Paired-sample t-test was performed to compare firing rates of the same sensillum when exposed to odor at different concentrations. The comparison was only performed between neighboring concentrations. Significance levels (0.05) were Bonferroni-corrected for multiple comparisons.

We performed one-sample t-test on behavioral data to identify concentrations with a mean behavior preference index significantly different from 0 (significance level = 0.05).

We tested the normality of data using the Jarque-Bera test.

### Data availability

The data used in this study can be made available on reasonable request to the authors.

## Acknowledgements

We thank Johannes Reisert (Monell Chemical Chemical Science Center), Vikas Bhandawat (Duke University) and members of Raman Lab for useful feedback on earlier versions of this manuscript. This research was supported by an Office of Naval Research grant (N00014-12-1-0089), and a NSF CAREER grant (#1453022) to B.R.

## Author Contributions

B.R. conceived the study. P.D. and H.R. performed the electrophysiology experiments with help from D.S. A.L, H.R., and D.S. collected the T-Maze datasets and H. R. conducted the geotaxis behavioral studies. H.R. did all the analysis and modeling portions of the work. B.R. and H.R. wrote the paper, incorporating inputs and comments from all authors. B.R. and Y.B. provided overall supervision.

---

**Figure S1.**
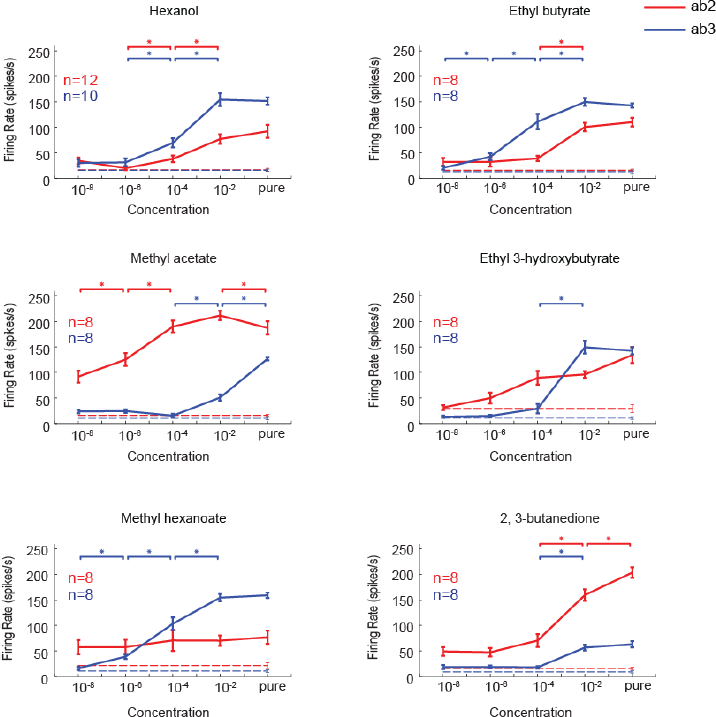
Dose response curves for both ab2 (red) and ab3 (blue) sensillum is shown for all six odorants used in this study. Asterisks indicate significant increase in firing rate compared with the neighboring lower concentration (p<0.05, paired *t*-tests). The dashed line indicates firing rate when presenting paraffin oil alone (i.e. the solvent used for diluting the odorants).

**Figure S2:**
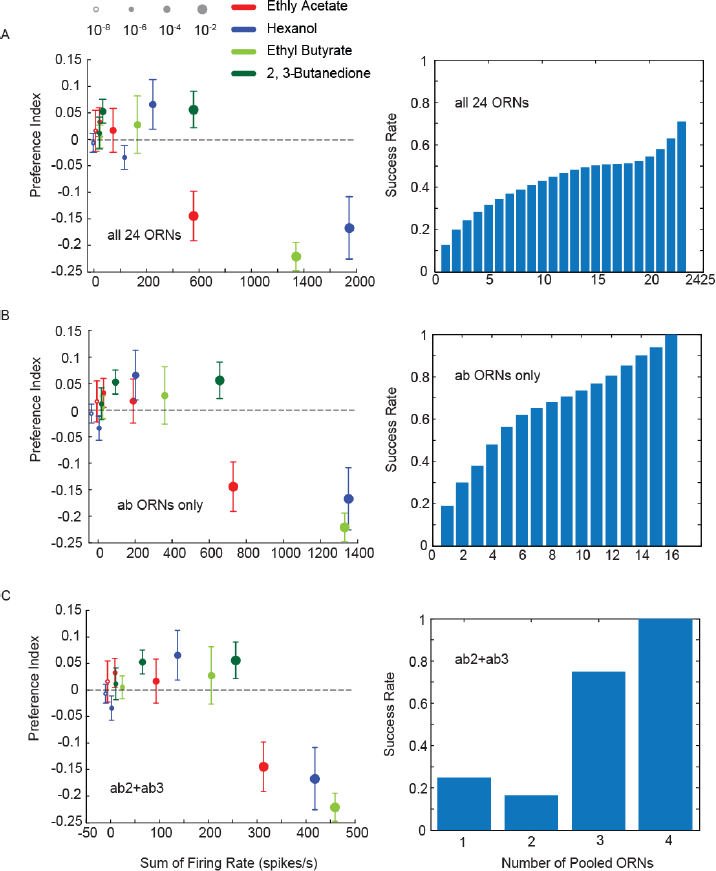
Independent validatiion of our results using a published dataset. (A) Similar plots as in Figure 6 but plotting the behavioral preference indices obtained in our T-maze experiments against cumulative spike counts of 24 different types of receptor neurons published in Hallem and Calrson (2006). Right panel reveals that monotonic increase in performance (i.e. correct recognition of the repulsive stimuli) as the number of neurons pooled for the analysis was systematically increased. Mean performance across different combinations of realizing a particular number of ORNs is shown along with SEM (i.e. 24 choose ‘n’ for any n ORN combination). (B) Similar plot as in panel a, but revealing prediction performance when selectively combining spiking activities of all ORNs housed in antennal basiconic type sensilla is shown. (C) Repeat of analysis in Figure 6b but using Hallem and Carlson (2006) data. Note that the analysis was limited to four odorants used in both our work and in the previous study.

